# A programmable and automated optical electrowetting-on-dielectric (oEWOD) driven platform for massively parallel and sequential processing of single cell assay operations

**DOI:** 10.1101/2024.03.19.585693

**Authors:** Lawrence G. Welch, Jasper Estranero, Panagiotis Tourlomousis, Robert C. R. Wootton, Valentin Radu, Carlos González-Fernández, Tim J. Puchtler, Claire M. Murzeau, Nele M. G. Dieckmann, Aya Shibahara, Brooke W. Longbottom, Clare E. Bryant, Emma L. Talbot

**Author notes:** These authors contributed equally (Lawrence G. Welch, Jasper Estranero, Panagiotis Tourlomousis).

## Abstract

Recently, there has been an increasing emphasis on single cell profiling for high-throughput screening workflows in drug discovery and life sciences research. However, the biology underpinning these screens is often complex and is insufficiently addressed by singleplex assay screens. Traditional single cell screening technologies have created powerful sets of ‘omic data that allow users to bioinformatically infer biological function, but have as of yet not empowered direct functional analysis at the level of each individual cell. Consequently, screening campaigns often require multiple secondary screens leading to laborious, time-consuming and expensive workflows in which attrition points may not be queried until late in the process. We describe a platform that harnesses droplet microfluidics and optical electrowetting-on-dielectric (oEWOD) to perform highly-controlled sequential and multiplexed single cell assays in massively parallelised workflows to enable complex cell profiling during screening. Soluble reagents or objects, such as cells or assay beads, are encapsulated into droplets of media in fluorous oil and are actively filtered based on size and optical features ensuring only desirable droplets (e.g. single cell droplets) are retained for analysis, thereby overcoming the Poisson probability distribution. Droplets are stored in an array on a temperature-controlled chip and the history of individual droplets is logged from the point of filter until completion of the workflow. On chip, droplets are subject to an automated and flexible suite of operations including the merging of sample droplets and the fluorescent acquisition of assay readouts to enable complex sequential assay workflows. To demonstrate the broad utility of the platform, we present examples of single-cell functional workflows for various applications such as antibody discovery, infectious disease, and cell and gene therapy.

## Introduction

Droplet microfluidics offers a range of properties that are highly advantageous to drug discovery and life sciences research.^1^ Fine control over temperature,^2^ cell confinement,^3^ and concentration gradients^4^ allow for high definition exploration of experimental space in a rapid and easy manner. Each droplet can be considered as a separate, isolated reaction or incubation vessel allowing massively increased experimental throughput with miniaturised cell and reagent consumption in a greatly reduced laboratory footprint. The dropletisation of cells into picolitre-scale volumes means secreted products rapidly accumulate to high concentrations enabling faster readouts for secretion-based assays at single-cell scale.^5^ Furthermore, inter-droplet crosstalk is eliminated by the use of fluorous continuous phases and stabilising surfactants.^6, 7^

Recently, the focus on high-throughput screening in drug discovery and the life sciences has shifted from bulk analyses to single cell profiling.^8^ However, the full depth of the biological complexity that is being queried in these screening campaigns is rarely fully addressed by single assay readouts alone. Conventional single cell screening methods usually offer powerful, but limited insights that may not constitute true functional analysis.^9^ As a result, these screening efforts often necessitate numerous additional secondary screens, making the process labour-intensive, lengthy, and costly. Consequently, key attrition points are not addressed until the late stages of workflows which further exacerbates costs and delays. Microfluidic systems to address singly encapsulated cells in droplets have been developed, but often suffer from limitations. For example, after encapsulation, droplets are manipulated using various force combinations within the extremely low Reynolds number environments typically found within microfluidic devices.^10^ Addressable arrays of confined fluid can be isolated using valve arrays^11^ and fabricated wells,^12^ as well as more sophisticated droplet traps where droplets are held in place by flow fields^13^ or Laplace pressure.^14^ Typically a droplet array will be a balance of payoffs. Passive arrays operating via flow fields or Laplace pressures are simple to use and straightforward to fabricate but operate solely under passive control, whereas valved systems are individually addressable but complex to manufacture and require complex control algorithms.

Digital microfluidic systems have been developed that allow the manipulation of droplets through electrowetting-on-dielectric (EWOD).^15-17^ Briefly, an electric field is applied between top and bottom plates of the chip, causing the droplet to deform due to the charge induced on the inner droplet surface. By defining the shape of the conductors between which the field is applied, this deformation can induce droplet motion, or hold droplets stationary against a flow field. Such systems allow a fine degree of control over the positioning of each droplet but the need for a high definition electrode array to actuate individual droplets in an addressable way limits the realistic array size accessible to the technique, with electrode sizes typically in the hundreds of μm.^18^ Using thin-film transistor (TFT) technology addressable arrays of up to 46000 electrodes have been reported, giving a realistic droplet capacity of around 2500.^19^

We describe a platform that harnesses droplet microfluidics and optical electrowetting-on-dielectric (oEWOD) to perform sequential and multiplexed single cell assays in massively parallelised workflows to enable complex cell profiling during screening (Fig 1.). In both oEWOD and EWOD the droplets are manipulated using a regional change in the contact angle between the droplets and the surface.^20^ In oEWOD the electrode shape is varied by using localised illumination to alter the conductivity of a photoactive layer built into the chip surface. This removes the need for complex pre-fabricated electrode arrays. In contrast, the projection of arbitrary light patterns is well developed, meaning the platform allows users to manipulate droplets with great flexibility and precision in a highly automated and parallel manner across the two dimensional space. The platform offers a range of standard operations that can be scripted to suit the user’s specific workflow requirements.

**Fig. 1.**
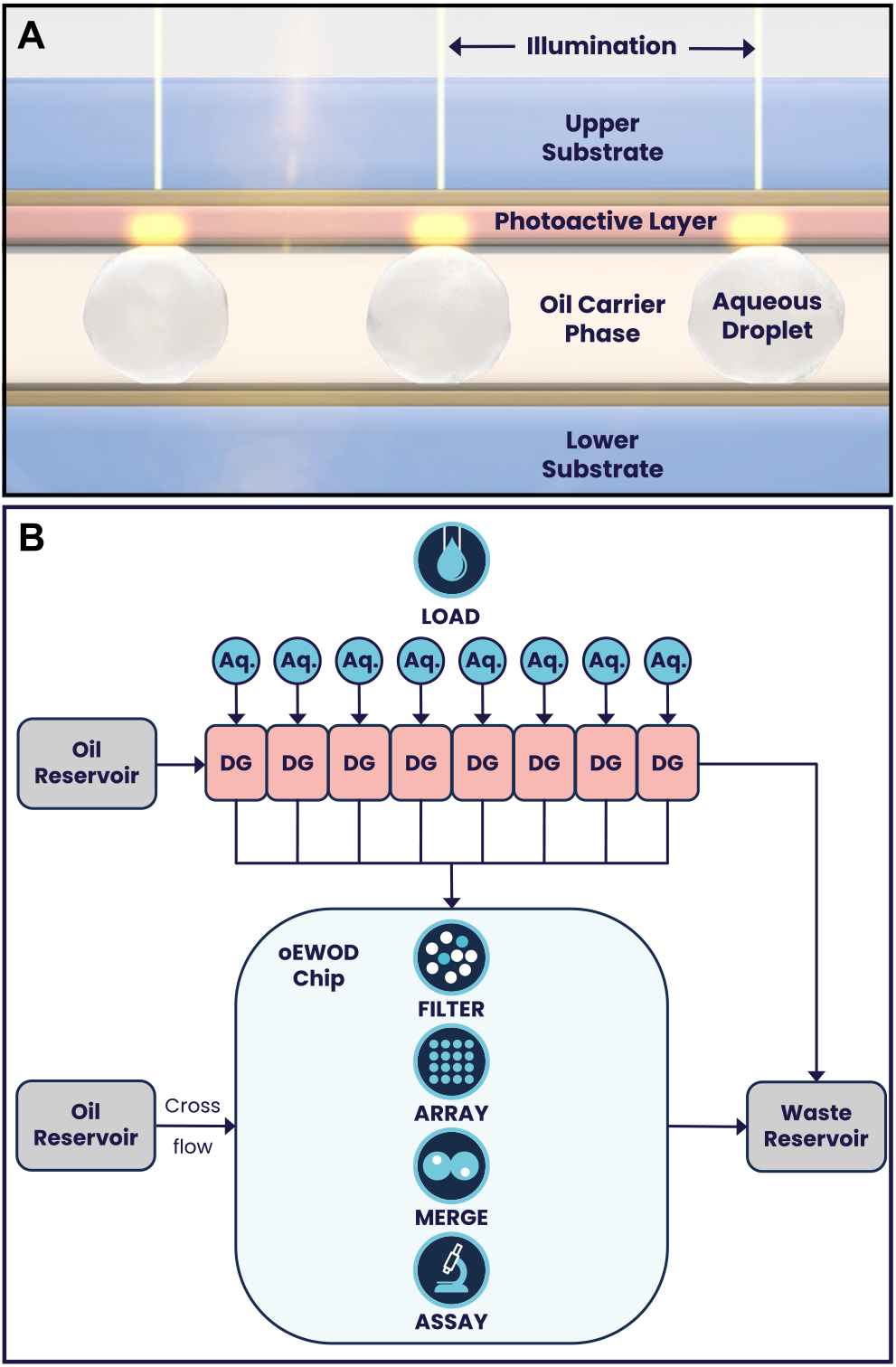
(A) An illustration depicting a cross-section of the oEWOD chip. Two planar substrates are separated, creating a cavity in which droplets and carrier-phase liquids are contained. Transparent conductive oxide layers in the substrates allow the application of a voltage across the chip. One substrate contains a photoactive layer, allowing the local electric field to be changed by the application of light. Thus droplet motion can be controlled by application of patterns of light onto the chip. (B) A topological diagram of the platform. Fluorous oil and up to 8 aqueous (Aq.) inputs are combined to form monodisperse droplets using droplet generator (DG) chips. These droplets are conveyed to the chip where they are filtered, undesirable droplets are removed, and the remainder formed into arrays ready for merger and acquisition events. The chip has the capability to cross flow oil in order to maintain optimum droplet conditions.

### Load and Filter

Objects in aqueous suspension, such as cells or beads, are encapsulated into picolitre-scale droplets (typically around 100-500 pL with diameters ranging from 60 to 100 μm). These droplets are created using low shear-stress droplet generators (DG) using a continuous phase of HFE7500 (NOVEC, 3M, Bracknell, UK) containing a block copolymer fluorosurfactant (Emulseo, Pessac, France) and are moved onto the temperature-controlled oEWOD device using pressure induced flow.^21^ On chip, the droplets are manipulated by oEWOD through the automated projection of individual light patterns, where individual points of light are termed sprites. Once assigned to a sprite, the history and assay performance of an individual droplet and its contents are logged throughout the workflow. Each droplet is assigned a unique ID, eliminating the need for a physical barcode. Under the control of light, droplets are conveyed across a detection window where they are visualised by a camera and are filtered in real-time based on droplet size, occupancy and content size. During encapsulation, the number of objects that occupy each droplet follows a Poisson probability distribution meaning that the input droplets are of mixed occupancy prior to filtering.^21-23^ Using a neural network algorithm optimized for image labelling, droplets of the correct size and correct occupancy (typically, but not exclusively, single occupancy) are actively identified and marked for retention. These target droplets are subsequently transported to an array and stored on chip pending further processing. Conversely, droplets with the incorrect size, content count, or content size are disqualified and sent to waste, ensuring the array only contains droplets with the content of interest.

### Merge

Many biological screening workflows involve the mixing of different assay reagents and/or objects in order to initiate a reaction or a cellular response. On the platform, this capability is enabled by the merge function. Merger of the droplets is effected by a pre-programmed motion sequence which brings droplets into contact and then oscillates them until the barrier to merger breaks down due to repositioning of the outer surfactant layer. The merged droplet is then treated as a new, single entity within the platform and the software retains the complete droplet history of parent droplets. Biological assays often require the combination of more than two mixtures. Furthermore, assays may be sensitive to the sequence and timing of component mixing. For example, one assay may require the mixing of multiple components at the same time while another may require the addition of a final component after a time delay such as after a baseline reading or after an incubation period. The flexible, programable nature of the platform enables sequential Merge workflows which go beyond two-way merge operations and which facilitate complex biological experimentation.

Combining Load, Filter and Merge with the ability to image fluorescent reporters in multiple channels, it is possible to design complex biological workflows and run them in a massively parallel manner. In this report, we initially use fluorescent beads to validate the core functional features of the platform. We subsequently illustrate the broad utility of the platform to life science research and drug discovery through demonstration of workflows spanning antibody discovery, infectious disease and cell and gene therapy.

## Material and Methods

### Cell culture and treatments

All cell culture reagents and treatments were supplied by Themo Fisher Scientific (UK) unless stated otherwise. All mammalian cells were cultured at 37°C in a humidified incubator with 5% CO_2_ until required for use. All cells were routinely screened for mycoplasma contamination. Freestyle CHO-S cells (Thermo Fisher Scientific, UK) were cultured in BalanCD CHO Growth A medium (Irvine Scientific, the Netherlands) supplemented with 8 mM L-glutamine in Erlenmeyer flasks in a shaking incubator at 125 rpm. Immortalised bone marrow-derived macrophages (BMDMs) and SK-N-AS (ATCC, CRL-2137) adherent cell lines were cultured in Dulbecco’s media with high glucose, 10% fetal bovine serum (FBS) and 1% penicillin-streptomycin (PS). Prior to loading on device, adherent cells (BMDMs and SK-N-AS) were dissociated from flasks in which they were thoroughly washed three times with Mg^2+^ and Ca^2+^ deficient PBS, and treated with trypsin for 5-10 minutes and neutralised in complete medium. Cells were pelleted at 250-300 x *g* for 5 minutes and resuspended in complete medium. Jurkat cells (ATCC, E6-1) were cultured in RPMI 1640 with GlutaMAX with 10% FBS and 1% PS and anti-TNFα hybridoma cells (ECACC, 2-179-E11) in RPMI 1640 with GlutaMAX with 10% FBS without PS. CD19+ B-cells (HemaCare, US) were cultured in B cell Excellerate medium with the CEllXVivo B cell expansion kit (R&D systems, UK). CD8+ T-cells (Cambridge Bioscience, UK) were thawed the day before the assay and were cultured on plates precoated with OKT3 antibody in Jurkat culture medium supplemented with 35 ng/mL IL-2. The NK cell line KHYG-1 (DSMZ, Germany) were cultured in Jurkat medium supplemented with 10 ng/mL IL-2. Overnight cultures derived from a single colony of GFP modified *Salmonella typhimurium* (L1344 strain)^24^ were prepared in LB Broth at 37°C with 200 rpm agitation. The working culture was prepared by inoculating 10% v/v of the overnight culture into fresh LB Broth followed by further agitation at 200 rpm for 2 hours at 37°C.

Where specified, cells were stained with CellTrace or CellTracker dyes in which cell suspensions were pelleted at 100-300 x *g* for 5 minutes, the supernatant removed and cells resuspended in serum-free basal media or phosphate-buffered saline (PBS). Cells were subsequently pelleted by centrifugation and resuspended in dye solutions (CellTracker DeepRed, 1-2 μM; CellTrace Violet, 2.5 μM; CellTracker Orange CMTMR, 5 μM; CellTrace Violet, 2.5 μM) and incubated at 37°C for 20-30 minutes. Cells were then pelleted by centrifugation at 100-300 x *g* for 5 minutes, the supernatant was removed and cells were washed in their respective complete medium.

### Bead Preparation

All beads were provided by Spherotech (US) unless stated otherwise. For the antibody discovery assay, 10-14 μm streptavidin-coated polystyrene beads were precoated with biotinylated human TNFα (R&D Systems, UK). Beads used for demonstration of the Load, Filter and Merge functions were 10-14 μm Nile Red Particles and 10-14 μm Yellow Particles. The cytokine capture bead for the infectious disease workflow was generated by conjugating an anti-mouse TNFα antibody (#6B8, BioLegend, UK) to 6.7 μm carboxyl polystyrene beads according to Spherotech guidelines.

### Input sample preparation

Prior to loading and after sample staining and washing, cells and beads were resuspended in complete medium with 12 or 20% OptiPrep (STEMCELL Technologies, UK) to limit object sedimentation during loading. For cell-based assays, the suspension solutions were adjusted to a final HEPES concentration of 25 mM. For the CHO Load and Filter viability experiment, cell viability was measured in suspension by trypan blue exclusion immediately before loading. Heat-kill controls were prepared in which 10% of cells were pre-stained with CellTrace Violet, heat-treated at 85°C for 5 minutes and allowed to cool to room temperature for 5 minutes. Cells were washed and mixed with non-heat-treated cells and the viability dye DRAQ7 (Abcam, UK) was added to a final concentration of 3 μM prior to loading as a mixed population. For the infectious disease workflow simulation, all input solutions were supplemented with 1 μg/mL of PE-conjugated rat anti-mouse TNFα antibody (MP6-XT22, BioLegend, UK).

### Microfluidic chip

oEWOD devices used for microdroplet manipulation were made consisting of two composite substrates – ‘active’ and ‘passive’ separated by a distance corresponding to less than a droplet diameter. Active substrates containing a photoconductive layer were comprised of: an optically transparent base substrate, an optically transparent conductor layer (70-250 nm), a photoactive layer (750-850 nm), dielectric layers deposited via ALD (<20 nm) and finally an anti-fouling, hydrophobic coating (1-15 nm). Passive layers were comprised of: an optically transparent base substrate, an optically transparent conductor layer (70-250 nm), dielectric layers deposited via ALD (<20 nm) and an anti-fouling, hydrophobic coating (1-15 nm). Electrical contacts were made by abrading the dielectric/photoactive layers in thin lines at the passive and active substrate ends to reveal the conductor layer and subsequent application of conductive paint.

### Droplet Generation

Droplets were generated using separate fluidic chips, fabricated out of polydimethylsiloxane (PDMS) by moulding,^25^ then coated in a hydrophobic material. In the emulsification chips the aqueous phase containing the intended contents of the droplets was flown into the carrier phase through a bank of nozzles with carefully designed geometry, producing droplets of a consistent size (diameter CV ∼ 10%) that were then transported to the microfluidic chip for use. The emulsification method was chosen as it represents a method that generates droplets with high-throughput, low variability and with low shear forces.

### Sample Load and Filtering

Droplet emulsions were transported into the microfluidic chip from the droplet generators under pressure. Upon entering the chip, they were spread-out by hydrodynamic forces and picked up by individual sprites, transferring their motion from fluid flow to oEWOD control. The droplets were then transported through a region imaged by a camera, allowing contents and sizes of droplets to be screened. Unwanted or unoccupied droplets were then moved under oEWOD control to a waste port to be disposed of, where hydrodynamic flow conveyed them to a waste reservoir, whilst those deemed acceptable were moved to a system-specified position on the chip for use.

### Optical Imaging

Sample imaging and read-out was performed using an *in-situ* infinity-corrected epifluorescence microscope. Illumination for both fluorescence and brightfield imaging consists of a broad-band white LED source passing through one of a selection of excitation filters (Semrock) before a multi-band dichroic beam-splitter. The light was then sent to the sample via a chosen objective. Fluorescence emission or reflected brightfield light was then collected using the same objective, passing through the multi-band dichroic and one of a selection of emission filters before being focussed onto an Andor Zyla 4.2 scientific CMOS (sCMOS) camera.

### Load and Filter Model

Prior to being stored in an array, droplets were traversed through a detection window in which a neural network classified the droplets and assigned a probability to their content number. Once a droplet reached the end of the window, a consensus decision was made based on all the individual probabilities at each time-point, and the droplet was discarded or sent to the array accordingly.

### Assay Signal Measurement

The droplet array on chip was imaged at user defined intervals. Whenever an image was saved, if there was a brightfield frame, the droplets were detected and their position and size was recorded. If the image was a fluorescence frame, the pixels were automatically segmented, and objects detected and profiled. From this process, information of their brightness, position and size was saved into a database that was accessible in real time.

## Results

### Load and Filter

To demonstrate the capability of the Load and Filter functions of the platform, 10 μm fluorescent polystyrene beads were encapsulated into 100 μm droplets (Fig. 2A; Supplementary Video 1). Under the control of oEWOD, bead-containing droplets of mixed occupancy were conveyed across the detection region where they were actively visualised and processed. Using a neural network optimized for image labelling, single occupancy droplets of the correct size (100 μm) were identified and marked-up in software register for retention. These target droplets were subsequently guided to an array and stored on chip pending further processing (Fig. 2C). In contrast; empty droplets, droplets containing multiple beads or incorrectly-sized droplets were disqualified and sent to waste. Waste droplets were routed so as to avoid array areas and minimise the risk of fouling. Furthermore, as the position of the droplets in the array is not defined by the physical chip structure (no fixed electrode array), this affords a flexibility in assay design. Arrays can be modified to meet specific assay requirements and can range in scale from tens of thousands of droplets for a meso-sized chip (covering an area of approximately 5.5 cm x 2.5 cm; as demonstrated by the array in Fig. 2B) to hundreds of thousands of droplets for larger chips.

**Fig. 2.**
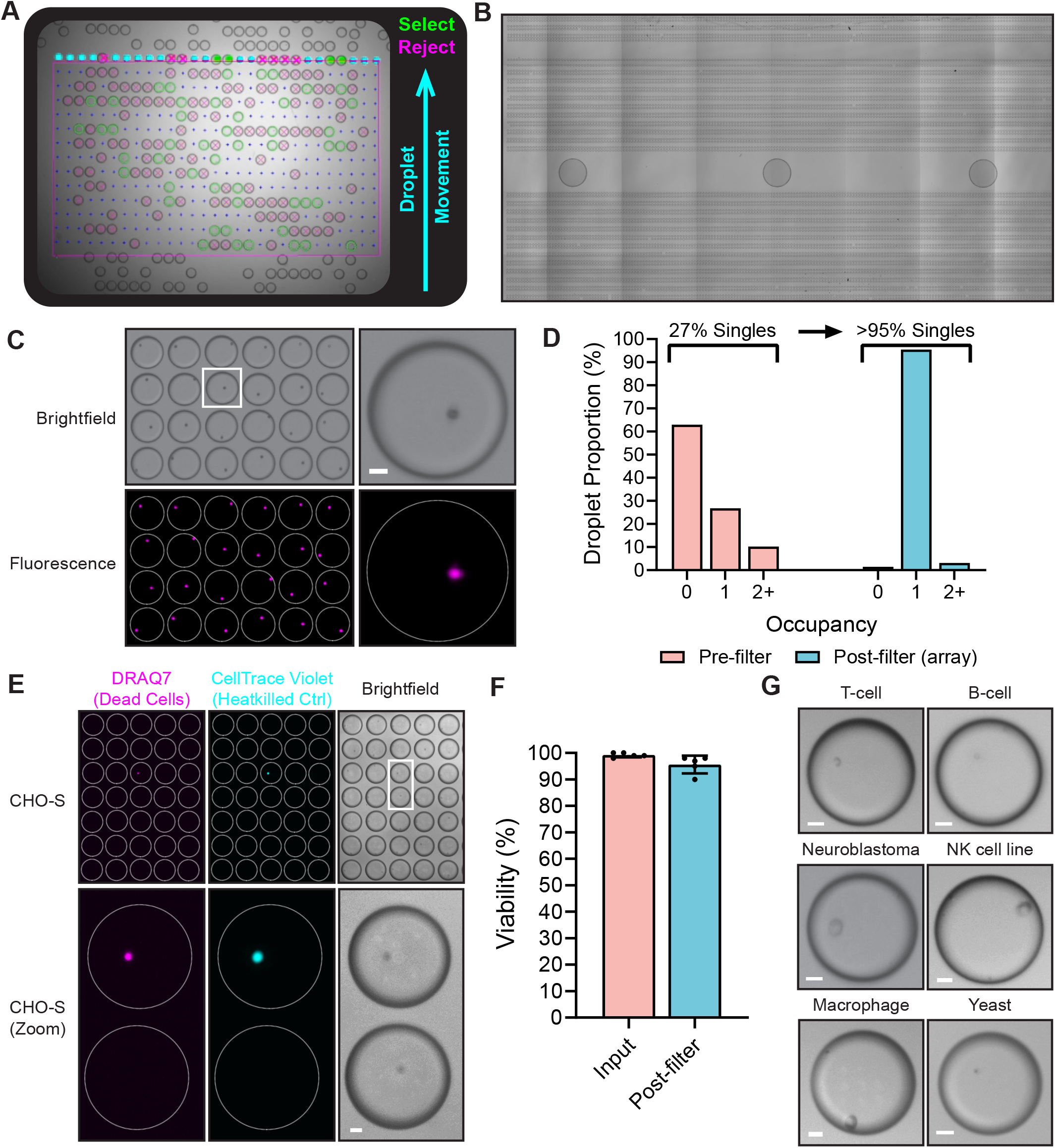
(A) A still image from a video showing the real-time droplet Filter function on the platform with annotations (see Supplementary Video 1); demonstrated using 10 μm fluorescent beads sorted for single occupancy in 100 μm droplets. Markup; blue arrow - direction of droplet movement, green – desirable droplets marked for retention in the array, magenta – undesirable droplets sent to waste. (B) A stitched image showing 12,932 aqueous droplets arrayed across a section of a chip after loading and filtering based on size (100 μm diameter). (C) A field of view of the array after the Load and Filter functions shown in (A), acquired using brightfield and fluorescence imaging (far-red, Nile Red). (D) The droplet object occupancy distribution before content filtering and after filtering when stored in an array on chip, as shown in (C). (E) The Lightcast platform can load and singulate cells in a gentle, non-toxic manner. Shown is a reduced field of view of CHO-S cells arrayed on device. Dead cells were identified with the viability dye DRAQ7. 10% of the input were heat-killed cells (pre-stained with CellTrace Violet) to confirm the efficacy of the viability dye. (F) Bar graph comparing the mean viabilities of CHO-S cells when in suspension prior to loading (input) and after Load and Filter in an on chip array (post-filter). Error bars, standard deviation; N = 5 (biological replicates). (G) Micrographs of a wide range of cell types which were successfully loaded and filtered for single occupancy on device. Videos and micrographs were captured using a 2x magnification objective in (A) and (B) and a 4x objective in (C), (E) and Scale bars, 20 μm. White rings indicate droplet outline.

Typically droplets are moved at a rate of 100-200 μm/s and for a load without content filter requirements (i.e. droplets containing solutions without objects) we loaded 12,932 droplets in under 20 minutes (Fig. 2B). However, when loading samples containing objects, the time taken to complete a Load and Filter may vary depending on various factors such as the droplet occupancy distribution (see below). The software allows users to define load parameters according to array target size and/or the time taken to load. This enables users to fine tune their on device workflow to meet the specific requirements of their application – whether the user has a time-sensitive workflow or they are prioritising assay throughput. In the example in Fig 2A and C; the sorting algorithm detected and sorted the target of 10,000 single bead droplets in 1 hour and 23 minutes – querying a total of 75,693 droplets in the process. During encapsulation, the number of beads that occupy each droplet is driven by Poissonian statistics and is influenced by object behaviour in suspension meaning that the input droplets are of mixed occupancy prior to filtering. As the droplets are passed through the filtration stage, multiple brightfield images are taken of the contents of each droplet, allowing the system to create a consensus evaluation of whether the droplets are empty, have single occupancy, or have multiple occupancy. Only those with single occupancy are moved to the experimental array, while others are diverted to waste, thus overcoming the Poisson probability distribution and enriching the desired downstream experimental targets. In the loading example provided above, 27% of droplets from the droplet generator contained a single bead, while 63% were empty and 10% contained more than one bead (Fig. 2D). Following filtering, >95% of the droplets enriched in the experimental array were single occupancy. This demonstrates the power of the Filter function in overcoming the limitations of the Poisson distribution to massively increase assay throughput.

Flow cytometers are a commonly used tool in cell biology research for single cell cloning,^26^ however, it is known that cells may be subject to considerable shear stresses during sorting which can have implications for cell health and performance.^27^ This is because cells experience strong shear in a viscous medium during sorting, however, in the droplet generation system we describe, no comparable acceleration is experienced. We calculate the wall shear stress in our fluorescence-activated cell sorting (FACS) system to be 2.99 Pa. In contrast, the predicted shear stresses experienced during dropletisation are approximately 20x lower, calculated as 0.16 Pa. To explore this, CHO-S cells were loaded onto the platform and filtered for single occupancy. Cell viability was monitored using a combination of trypan blue exclusion and a fluorescent viability stain in which viability was measured for the input suspension (before loading) and after sorting into an array on chip (Fig. 2E). Across several loads, mean CHO cell viability remained over 95% before and after sorting suggesting that the Load and Filter function is indeed gentle and non-detrimental to cell viability (Fig. 2F). Beyond CHO cells, the Load and Filter function was applied to a wide range of different cell types (Fig. 2G) including primary cells (T and B cells), adherent cells (macrophages; SK-N-AS, neuroblastoma cell line), an immortalised suspension cell line (KHYG-1, NK cell line) and microbes (yeast). In summary, the Load and Filter functions are capable of singulating a broad range of cell types in a rapid, high-throughput and gentle manner.

### Merge

To demonstrate the Merge function, two bead types with different fluorescence profiles were sequentially loaded and filtered for single occupancy in 100 μm droplets (Fig. 3A). The droplets were arrayed in an interdigitated fashion meaning the droplet rows alternated between droplets containing 10 μm yellow fluorescent beads (Light Yellow) and those containing 10 μm far-red fluorescent beads (Nile Red). The interdigitated bead array can be seen in Fig. 3B. Arrayed droplets were subject to brightfield and fluorescent acquisitions on chip prior to the merging of neighbouring droplets under the orchestrated control of oEWOD (Supplementary Video 2). Droplets were merged in a parallel fashion as confirmed by a post-merge acquisition of the arrays. The ability to singulate and combine two objects with high precision and at scale is a powerful feature of the platform. Without Load, Filter and Merge; the probability of two different single objects being combined in the same droplet is very low as the Poissonian probabilities are compounded per object type. For example, the proportion of single occupancy droplets was 13.2% for Nile Red beads and 13.3% for Light Yellow beads prior to Filter. Without Filter, in this particular experiment there would only be a 1.8% chance of successfully combining one bead of each type into the same droplet as the single occupancy probabilities for each load are multiplied. With Filter, the proportion of single occupancy droplets in the array was increased to 96.9% for Nile Red beads and 95.1% for Light Yellow beads giving a compound probability of 92.1%; a figure which is considerably higher than comparable metrics associated with other droplet microfluidics devices.^28-30^

**Fig. 3.**
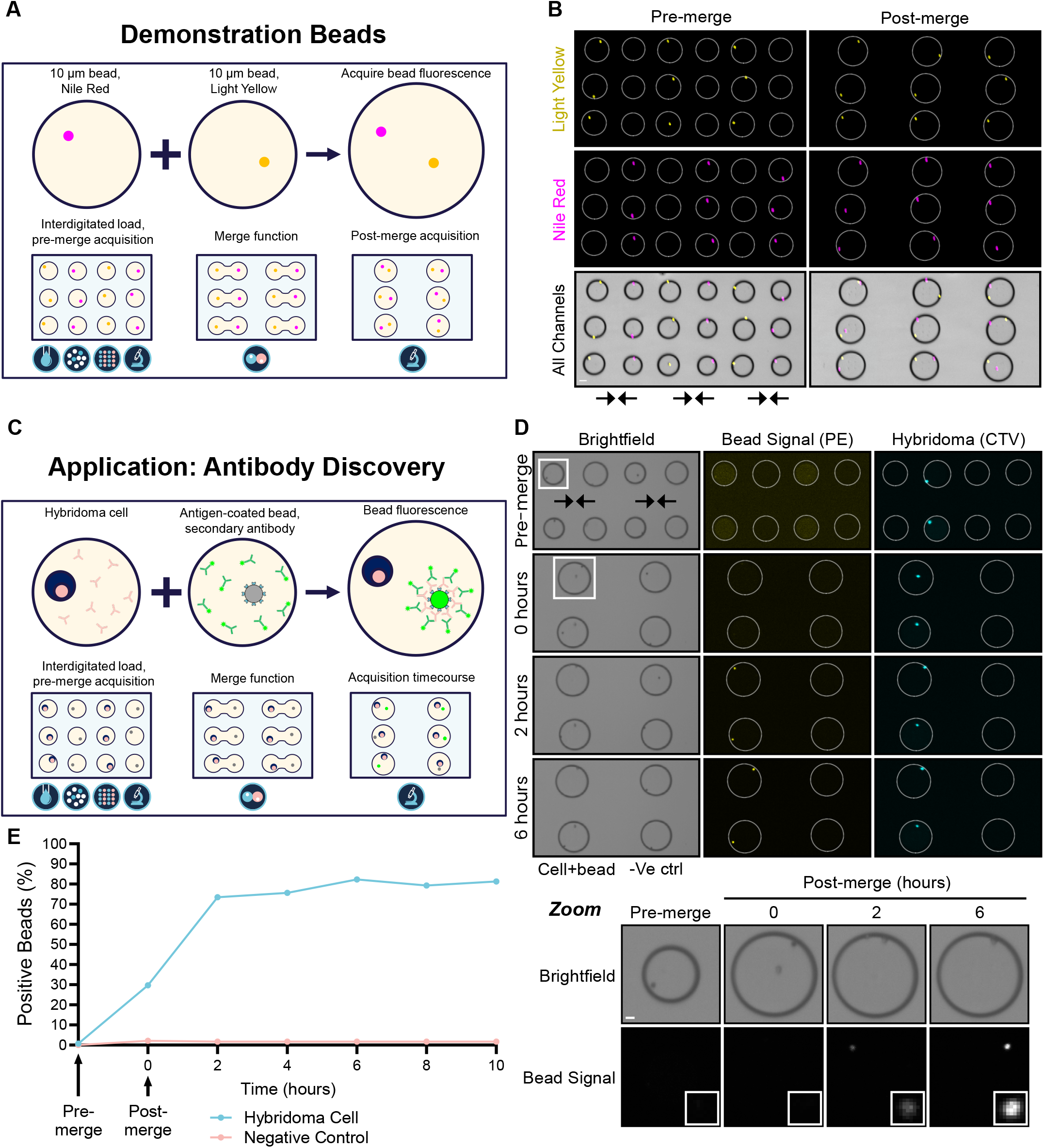
(A) A schematic of the Merge function demonstrated using two differentially fluorescent beads. Two bead suspensions containing 10 μm beads fluorescent in red/far-red (Nile Red) or in yellow were separately dropletised, loaded, filtered for single occupancy and moved into arrays on the platform in an interdigitated fashion. During the preprogrammed Merge function, droplet pairs were brought together and fused under the control of oEWOD (see Supplementary Video 2). (B) Fluorescence and brightfield micrographs of bead-containing droplets before and after merge on chip. Shown is a small section of the array. Arrows indicate direction of merge. Scale bar, 50 μm; 2x magnification. White rings indicate droplet outline. (C) A schematic illustrating the application of the Merge function to an antibody discovery workflow. Pooled hybridoma clones are singulated, merged with a single antigen-coated bead and a fluorescent secondary antibody and are imaged over time. Hybridoma clones producing antibodies specific to the antigen of interest will cause the fluorescent signal to coalesce on the bead. (D) Micrographs of the workflow in C using TNFα-coated beads and a hybridoma clone producing antibodies specific to TNFα (cells stained with CellTrace Violet [CTV]). Shown is a small section of the array. The timecourse refers to hours after merge. Screening reactions are shown adjacent to negative control reactions in which beads were merged with empty droplets (just media, no cells). White box, inset; zoom of a cell-containing droplet showing bead signal over time. Scale bar, 20 μm; 2x magnification. (E) The percentage of fluorescent beads detected over time from the experiment shown in (D). Representative of 3 biological replicates.

To illustrate the value of the platform to life sciences research and development, a workflow was designed for a primary antibody discovery screen (Fig. 3C). The aim of the workflow is to screen for rare cell clones (e.g. hybridoma or B-cells) that produce antibodies that are specific to an antigen of interest such as a validated drug target. In the workflow, the different cell clones are encapsulated, filtered for single occupancy and merged with droplets containing a fluorescently-labelled secondary antibody and single beads coated with the antigen of interest. Secreted antibodies that are specific to the antigen of interest will bind to the antigen-coated bead. This will in turn recruit the secondary antibody causing a fluorescent signal to coalesce on the bead, thereby signifying the clone is a positive hit. In contrast, in merged droplets without cells or with cells producing non-specific antibody clones, the beads remain non-fluorescent. To test the workflow, a validated hybridoma clone was selected that is known to produce an antibody specific to the human cytokine TNFα. Beads were precoated with TNFα and were resuspended in media containing a PE-conjugated anti-mouse secondary antibody. The bead mixture was loaded, filtered for single occupancy and merged with single anti-TNFα hybridoma cells (stained with CellTrace Violet) or droplets lacking cells as a negative control (Fig. 3D). Brightfield and fluorescent images were captured before merge and for several hours after merge in order to track the changes in bead brightness over time. As expected, there was no change in bead intensity for the negative control (Fig. 3E). In contrast, an increase in bead brightness was detectable for droplets containing hybridoma cells immediately after merge (within 15 minutes) demonstrating the sensitivity and speed of the assay at single cell resolution. Furthermore, the bead intensity continued to increase for at least 6 hours after merge, highlighting that assays on the platform can monitor changes over time unlike many traditional endpoint immunoassay formats such as ELISAs.

### Sequential Merge

As an illustration of a sequential merge, three distinct input samples were sequentially loaded, filtered and then arrayed in an interdigitated fashion with relation to one another (Fig. 4A). The aim of the experiment was to simulate a potential sequential merge sequence applicable to infectious disease research. In this workflow, one droplet population contained single macrophages, another contained single immunoassay cytokine capture beads and the final droplet contained the bacterium *Salmonella typhimurium*. In the presence of secreted cytokine, a fluorescently-labelled detection antibody is recruited to the bead in an immunoassay sandwich complex resulting in a fluorescent bead signal. The initial merge of the bead and cell droplets enables the measurement of the baseline cytokine secretion level of the macrophage. The subsequent merge of the third droplet population serves to assess the cytokine secretion rate and viability of single macrophages after challenge with *Salmonella*, with reference to baseline levels. During the on chip simulation of the workflow; cytokine beads, macrophages and *Salmonella* were successfully encapsulated, filtered and sequentially merged with precision in an automated parallelised fashion (Fig. 4B, Supplementary Video 3).

**Fig. 4.**
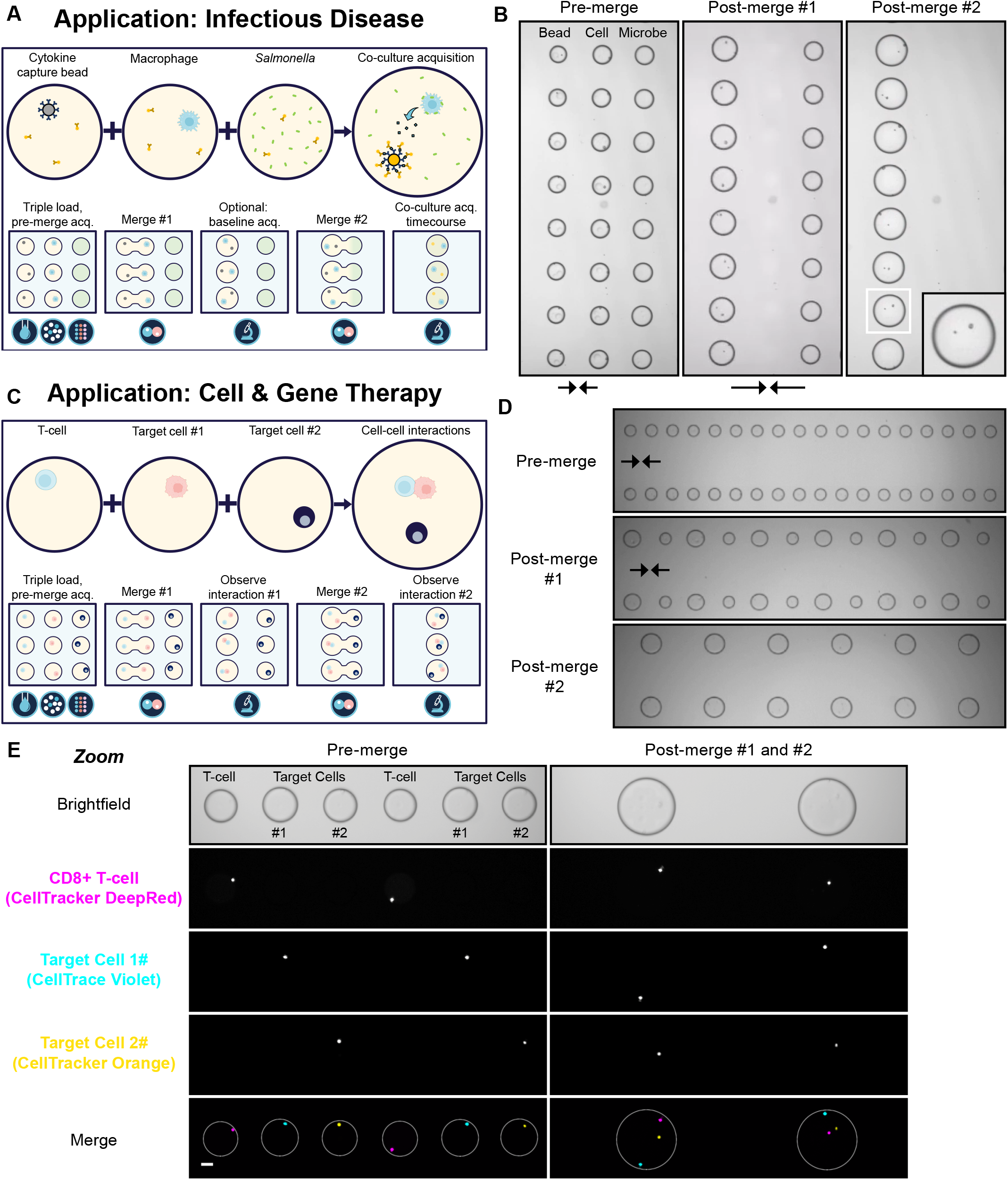
(A) A schematic of a sequential merge workflow applicable to infectious disease research. Droplets containing single cytokine capture beads and macrophages are merged and baseline cytokine secretion levels are measured using a fluorescently-labelled cytokine-specific detection antibody. The bead and cell droplets are subsequently merged with GFP-positive *Salmonella* droplets and changes in cytokine secretion and viability are tracked over time. (B) Video stills from a proof of concept simulation of the workflow illustrated in (A), arrows indicate direction of merge (see Supplementary Video 3). Zoom of a single droplet shown inset. Video captured with a 2x objective. (C) A schematic of a sequential merge workflow applicable to cell and gene therapy. Droplets containing single primary T-cells undergo two rounds of merging with droplets containing a single target cell. (D) Video stills from a proof-of-concept simulation of the workflow illustrated in (C), captured as in (B), (see Supplementary Video 4). (E) Micrographs of the workflow illustrated in (C) at 4x magnification. Scale bar, 50 μm. Differentially stained Jurkat cells were used as target cells for illustrative purposes.White rings indicate droplet outline. All micrographs show a small section of the total array.

To further illustrate the broad utility of the platform, we carried out of a proof-of-concept simulation of a workflow applicable to cell and gene therapy development. In this workflow, three different cell populations; an effector cell (e.g. a T-cell or NK cell) and two different target cell populations were combined through a sequential merge (Fig. 4C). The proposed workflow seeks to assess the ability of the effector cell to engage with, and ultimately kill, the target cells in an immune cell killing assay. The target cells may represent two of the same cell type, such as in a workflow designed to assess the serial killer capacity of effector cell clones. Alternatively, the target cells may vary, such as in a workflow which interrogates the specificity and/or range of the effector cell killing activity. In the proof-of-concept simulation, CD8+ primary T-cells (stained with CellTracker Deep Red) served as the effector cell population and two differentially-labelled Jurkat cells (stained with CellTrace Violet or CellTracker Orange) served as the target cell populations (Fig. 4D,E; Supplementary Video 4). As for the infectious disease workflow, the effector and target cell populations were successfully encapsulated, filtered and sequentially merged in synchrony on chip.

This workflow demonstrates that multiple different cell types can be singulated and combined with high precision and high parallelisation using the platform. Notably, the ability to overcome the limitations of the Poisson distribution becomes more powerful when attempting to combine more than two single objects in sequential merges. The single occupancy probability of T-cells and each Jurkat load was 28.1%, 26.9% and 26.7% respectively, before Filter. Therefore, after compounding the probabilities, the likelihood of combining one of each cell type into the same droplet is only 2.0% without Filter. With a typical Filter efficiency of 95%, in this example, the compound probability of correctly combining the three cell types into one droplet increases to >85%, an enrichment factor of >40. The single occupancy probabilities for each cell type were >25% before Filter as we increased cell densities in the input suspensions in order to minimise loading times. For other techniques without Filter, the input cell density is often much lower to minimise the incidence of multiple occupancy droplets (discussed below).^31^ In these instances, then the enrichment factor from the Filter function would be much greater. For example, were the probability of single occupancy <10% for the three cell populations discussed, then the likelihood of correctly co-encapsulating the three cell types may be as low as 0.1% and the enrichment factor offered by the Filter function could be as high as ∼850. With this unique and powerful capability to overcome the Poisson distribution, many previously impractical workflows in cell and gene therapy, and indeed beyond, can be made possible.

## Discussion

We have demonstrated the power of the Filter function for a variety of workflows including cells and beads where the desired outcome is to deterministically co-encapsulate the cells or beads in single droplets with high fidelity. The Poisson distribution issue is problematic for most other droplet microfluidic technologies (see review by Collins et al.)^23^ and this burden is further compounded when looking to co-encapsulate multiple different objects with precision. As a result, mitigation of the Poisson distribution issue has been the focus of intense development within the droplet microfluidics community.^31, 32^ This is critical as, for example, the performance of many traditional single cell ‘omic technologies is hampered by the contamination of the sample with droplets containing more than one cell.^31^ These technologies are unable to directly overcome the limitations of the Poisson distribution so users are required to massively reduce the density of their input suspension to make the proportion of droplets containing multiple cells negligible. This alteration in the distribution is at the cost of the proportion of desirable single cell droplets as the vast majority of the droplet population is likely to be empty. Consequently, these systems are required to screen a far higher volume of droplets in order to query the necessary number of single cells. In contrast, we have shown that the Filter function allows the platform to handle a range of different input object densities. To optimise our loading times, we generally use higher object input densities resulting in a higher proportion of single and multiple occupancy droplets pre-filter as the platform can efficiently remove undesirable droplets during Filter.

Another key defining difference from traditional single cell ‘omics technologies is that the platform does not require complex downstream deconvolution. As all of the images, the assay readouts and the history of each individual droplet is tracked from the point of Filter to the end of the workflow; there is no requirement for DNA barcoding, downstream sequencing or complex bioinformatic analysis for the core functionality of the platform. Instead, as the platform scans droplets that are stored and logged in the array, it can query true biological function in real-time, cell by cell and at scale. Furthermore, although the vast majority of undesirable droplets is eliminated by the Filter function, by imaging and assaying the arrayed droplets the platform has a second layer of quality control. The very small proportion (<5%) of mis-filtered empty or multiple occupancy droplets in the array can be belatedly disqualified or caveated during analysis of the array. Combined with this unparalleled double layer of stringency and the ability to assess each droplet individually in real-time, the negative impact of multiple occupancy droplets is negligible for workflows on the platform, unlike traditional technologies.

## Conclusions

We have described a novel platform that combines droplet microfluidics, oEWOD and complex software logic to enable the manipulation and assay of single cells with unprecedented throughput and complexity. We have demonstrated the unique power of the Filter function to overcome the Poisson distribution and enrich for droplets of the desired occupancy to massively increase assay throughput by optimising chip space. The flexible nature of oEWOD manipulation enables a suite of workflow modules such as Merge; which, when combined with Filter, enables complex sequential assay workflows involving multiple merge events. Droplets are stored in an addressable array on chip and their individual history is logged throughout the workflow allowing the biological function of single cells to be queried in real-time, cell by cell. Finally, we demonstrated the flexibility and broad applicability of the platform using bead-based models and with single-cell functional workflows spanning antibody discovery, infectious diseases, and cell and gene therapy. We believe the addition of deep functional profiling at single cell resolution and at scale provides a route to a more complete understanding of the underlying complexity of biology.

## Author Contributions

ELT and CEB conceived the study and reviewed the manuscript. LGW, RCRW, CGF, TJP and BWL wrote the manuscript. LGW, JGE, PT, VR, CMM, AS and NMGD collected the data.

## Acknowledgements

We thank Kathrin Herbst, Thomas Isaac, Matthew Wright, Scott Brouilette, Simon Margerison, Paul Steinberg, Jonathan Didier and Paul Loeffen for comments on the manuscript. We thank Nicole Lederer, Derya Emin, Nele Dieckmann, Joe Moore, Marina Efstratiou, Sion Harlow, Elisa Salis, Patricia Veiga Crespo, Leslie Oppon and Drew Geere with help with data collection.

